## Supplementary data for "RNA-Seq is not required to determine stable reference genes for qPCR normalization"

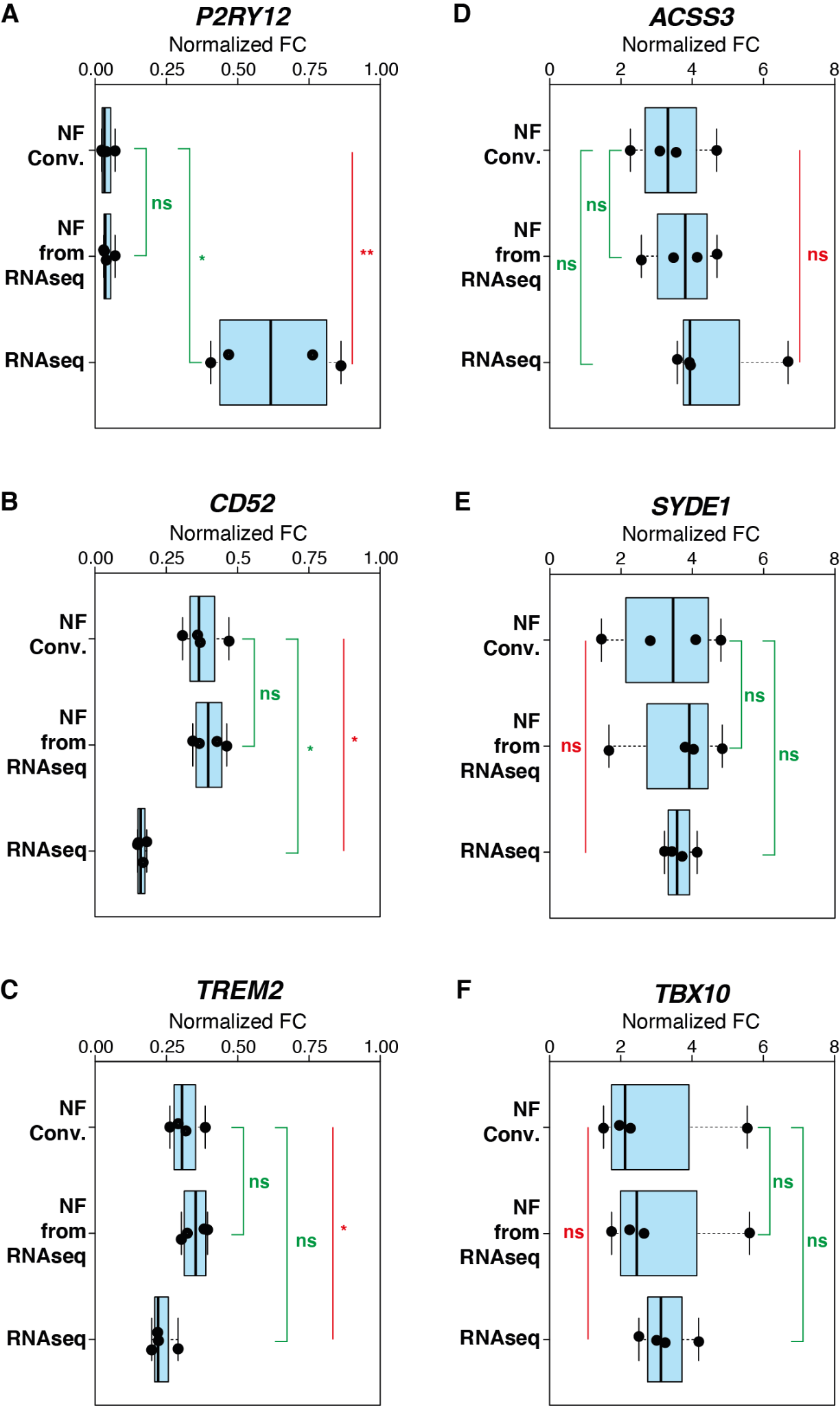

**Supplementary Figure 1:** Comparison of fold changes in TREM2 KO group among the three analysed methods (qPCR FC computed using NF Conv and NF RNA-Seq and finally RNA-Seq).

Non-parametric ANOVA was performed using Kruskal Wallis test (red line comparing the three groups). Multiple comparisons were performed using the Dunn’s post hoc test using the NF conv. group as the control condition (green lines). *The alpha value was set at 0.05 and P values are annotated as follows: * P<0.05, ** P<0.01, *** P<0.001.*

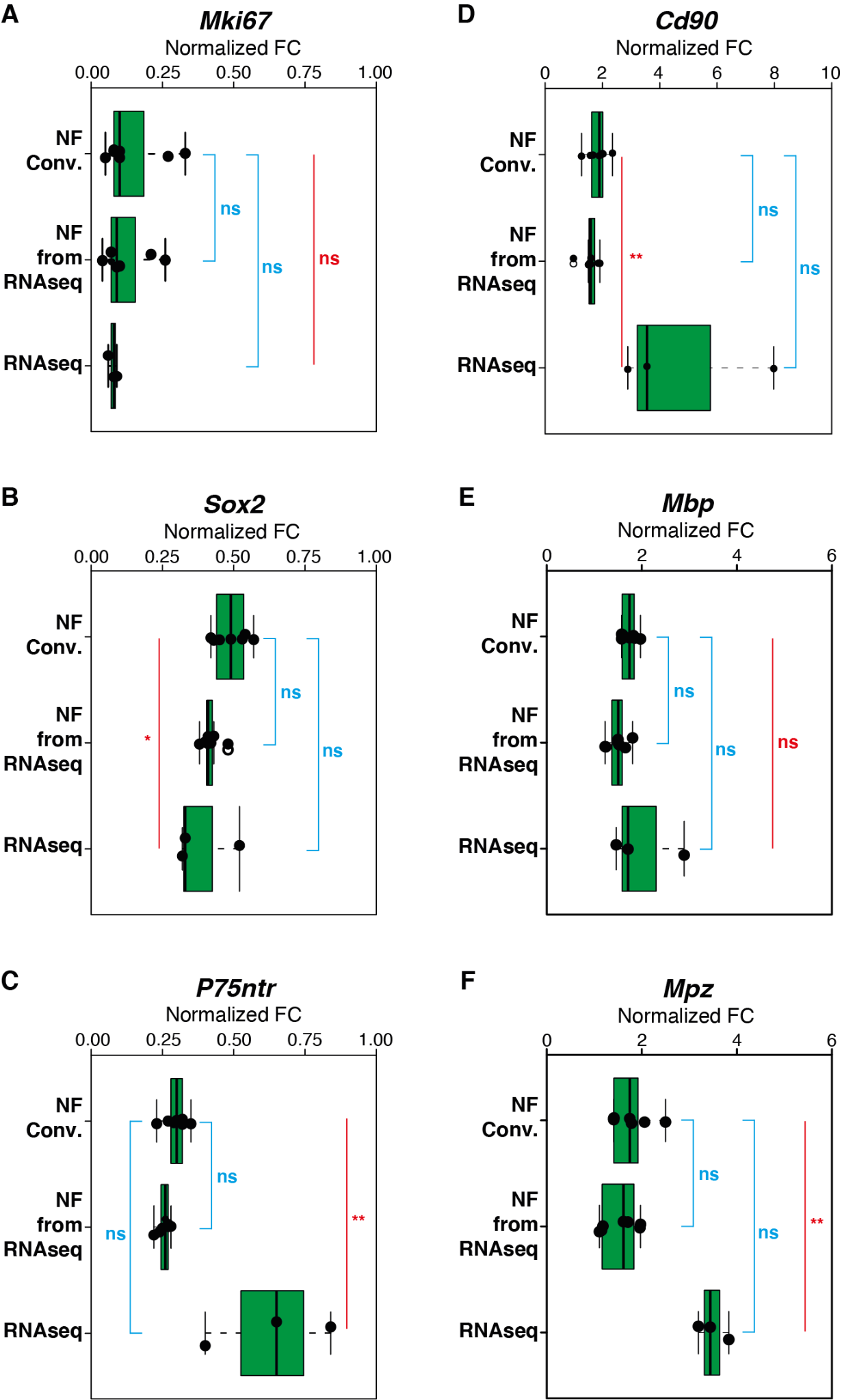

**Supplementary Figure 2:** Comparison of fold changes in TREM2 KO group among the three analysed methods (qPCR FC computed using NF Conv and NF RNA-Seq and finally RNA-Seq). Non-parametric ANOVA was performed using Kruskal Wallis test (red line comparing the three groups). Multiple comparisons were performed using the Dunn’s post hoc test using the NF conv. group as the control condition (Blue lines). *The alpha value was set at 0.05 and P values are annotated as follows: * P<0.05, ** P<0.01, *** P<0.001.*

**Supplementary Table 1. Reference genes chosen from RNA-Seq data WT vs TREM2KO**

| **ENSEMBL ID** | **Gene Name** | **baseMean** | **log2FC** | **pValue** | **pAdj** | **CVfromDisp** |
| --- | --- | --- | --- | --- | --- | --- |
| ENSG00000169714 | CNBP | 14427.26 | 0.05 | 0.68 | 0.84 | 12.41 |
| ENSG00000138279 | ANXA7 | 6823.55 | 0.09 | 0.43 | 0.66 | 11.66 |
| ENSG00000198231 | DDX42 | 6558.35 | -0.01 | 0.95 | 0.98 | 12.44 |
| ENSG00000111667 | USP5 | 5356.63 | -0.03 | 0.82 | 0.92 | 11.50 |
| ENSG00000204569 | PPP1R10 | 4105.63 | 0.10 | 0.42 | 0.66 | 11.84 |
| ENSG00000137177 | KIF13A | 3615.09 | -0.02 | 0.85 | 0.94 | 12.32 |
| ENSG00000164916 | FOXK1 | 2502.17 | -0.04 | 0.73 | 0.87 | 11.56 |
| ENSG00000149089 | APIP | 1996.63 | 0.04 | 0.72 | 0.87 | 12.00 |
| ENSG00000082258 | CCNT2 | 1652.96 | 0.05 | 0.72 | 0.87 | 12.22 |
| ENSG00000116221 | MRPL37 | 1521.71 | 0.04 | 0.76 | 0.89 | 12.06 |

**Supplementary Table 2. Reference genes chosen from RNA-Seq data P3 vs P21 Sciatic nerves**

| **ENSEMBL ID** | **Gene Name** | **baseMean** | **log2FC** | **pValue** | **pAdj** | **CVfromDisp** |
| --- | --- | --- | --- | --- | --- | --- |
| ENSMUSG00000020485 | Supt4a | 878.49 | 0.03 | 0.81 | 0.88 | 12.93 |
| ENSMUSG00000031513 | Leprotl1 | 964.45 | -0.01 | 0.94 | 0.96 | 12.80 |
| ENSMUSG00000032826 | Ank2 | 966.76 | 0.04 | 0.78 | 0.86 | 12.57 |
| ENSMUSG00000028581 | Laptm5 | 1020.98 | -0.03 | 0.86 | 0.92 | 12.89 |
| ENSMUSG00000031782 | Coq9 | 1049.41 | -0.06 | 0.68 | 0.79 | 11.99 |
| ENSMUSG00000030357 | Fkbp4 | 1095.09 | 0.08 | 0.57 | 0.71 | 12.76 |
| ENSMUSG00000028161 | Ppp3ca | 1937.16 | -0.05 | 0.74 | 0.84 | 12.76 |
| ENSMUSG00000033916 | Chmp2a | 3261.45 | 0.08 | 0.56 | 0.70 | 12.26 |
| ENSMUSG00000019899 | Lama2 | 5070.82 | 0.00 | 0.99 | 1.00 | 12.25 |
| ENSMUSG00000028452 | Vcp | 6228.91 | -0.10 | 0.48 | 0.63 | 12.34 |
